# Stability and sensitivity of structural connectomes: effect of thresholding and filtering and demonstration in neurodegeneration

**DOI:** 10.1101/416826

**Authors:** Peter McColgan, Tessel Blom, Geraint Rees, Kiran K Seunarine, Sarah Gregory, Eileanoir Johnson, Alexandra Durr, Raymund AC Roos, Rachael I Scahill, Chris A Clark, Sarah J Tabrizi, Adeel Razi, the Track-HD Investigators

## Abstract

Structural connectomes derived using diffusion tractography are increasingly used to investigate white matter connectivity in neurological diseases. However inherent biases in diffusion tractography algorithms may lead to both false negatives and false positives in connectome construction. A range of graph thresholding approaches and more recently several streamline filtering algorithms have been developed to address these issues. However there is no consensus in the literature regarding the best available approach. Using a cohort of Huntington’s disease patients and healthy controls we compared the effect of several graph thresholding strategies: proportional, absolute, consensus and consistency thresholding, with and without streamline filtering, using Spherical Deconvolution Informed Filtering of tractograms (SIFT2) algorithm. We examined the effect of thresholding strategies on the stability of graph theory metrics and the sensitivity of these measures in neurodegeneration. We show that while a number of graph thresholding procedures result in stable metrics across thresholds, the detection of group differences is highly variable. We also showed that the application of streamline filtering using SIFT2 resultes in better detection of group differences and stronger clinical correlations. We therefore conclude that the application of SIFT2 streamline filtering without graph thresholding may be sufficient for structural connectome construction.

## Introduction

Graph theory is a mathematical framework that can be used to study the organization of structural and functional brain networks. Networks are defined by a collection of nodes (brain regions) and links or edges between nodes. For structural connectivity, the edges represent white matter connections. Graph metrics can characterize one or several aspects of global or regional brain connectivity. Regional metrics can quantify the influence of a specific brain region in the network, while global metrics can provide information about the level of integration and segregation across the whole network (Rubinov and Sporns, 2010). Graph theory has been used to characterise abnormal structural connectivity in a range of neurodegenerative diseases including Alzheimer’s disease (Lo et al., 2010), frontotemporal dementia (Mandelli et al., 2016), amyotrophic lateral sclerosis (Verstraete et al., 2010) and Huntington’s disease (McColgan et al., 2015).

Network measures can be influenced by factors such as the choice of the brain parcellation scale, tractography algorithm, weighting scheme, thresholding approach and streamline filtering (Bullmore and Sporns, 2009; Qi et al., 2015; Smith et al., 2013). Tractography using MR diffusion imaging is a method that measures in-vivo anatomical connectivity by tracing the white matter fiber tracts. Deterministic tractography is a method based on the local fibre orientation derived from the principal directions of the diffusion tensor (Descoteaux et al., 2009). The inherent limitation of this technique is the inability to resolve crossing fibres leading to many false negatives in connectome reconstruction. This led to the development of probabilistic tractography based on the probability density distribution of the local fibre orientation (Behrens et al., 2007; Behrens et al., 2003). While this method is much more effective at resolving crossing fibres, it results in much denser connectomes and thus more false positives (Zalesky et al., 2016).

Graph thresholding is an effective and widely used strategy to remove the false positives created by the probabilistic approach (Achard and Bullmore, 2007; Rubinov and Sporns, 2010). As the name suggests, graph thresholding entails applying a quantitative threshold below which the connections are removed (by setting them to zero in the adjacency matrix) from further consideration. This procedure is useful on two accounts: firstly it helps to remove the spurious connections (the false positives discussed above) and secondly by inducing the sparsity that helps minimize the multiple comparison problem. This is potentially important because the metrics calculated from the ensuing sparse graphs are sensitive to the amount and the method of thresholding (Simpson et al., 2013). There are multiple thresholding approaches described in the literature - for example absolute, proportional, consensus and consistency thresholding. However currently there is no agreement in the literature regarding the best practice for threshold implementation (Qi et al., 2015). An absolute threshold defines a value, below which connections are removed (Figure 1, top row) (Daianu et al., 2015; Drakesmith et al., 2015; Li et al., 2016). A relative threshold retains a defined proportion of the strongest connections in the network (Figure 1, bottom row) (Mueller et al., 2015; Yao et al., 2010). Consensus thresholding retains only the connections present in a defined percentage of the group (McColgan et al., 2015; van den Heuvel and Sporns, 2011; van den Heuvel et al., 2013) (Figure 2). More recently, consistency thresholding has been proposed, whereby the most consistent connections across a group, as defined by coefficient of variation, are retained. Graph metrics may also be computed over a range of threshold values (Bai et al., 2012; Bassett et al., 2008). This is particularly common in the context of proportional thresholding. However, this can lead to false conclusions if the network measures are not stable across the range of thresholds (Scheinost et al., 2012). Previous work by (Garrison et al., 2015) has demonstrated the instability of absolute and proportional thresholding techniques in resting state fMRI functional brain networks.

**Figure 1.**
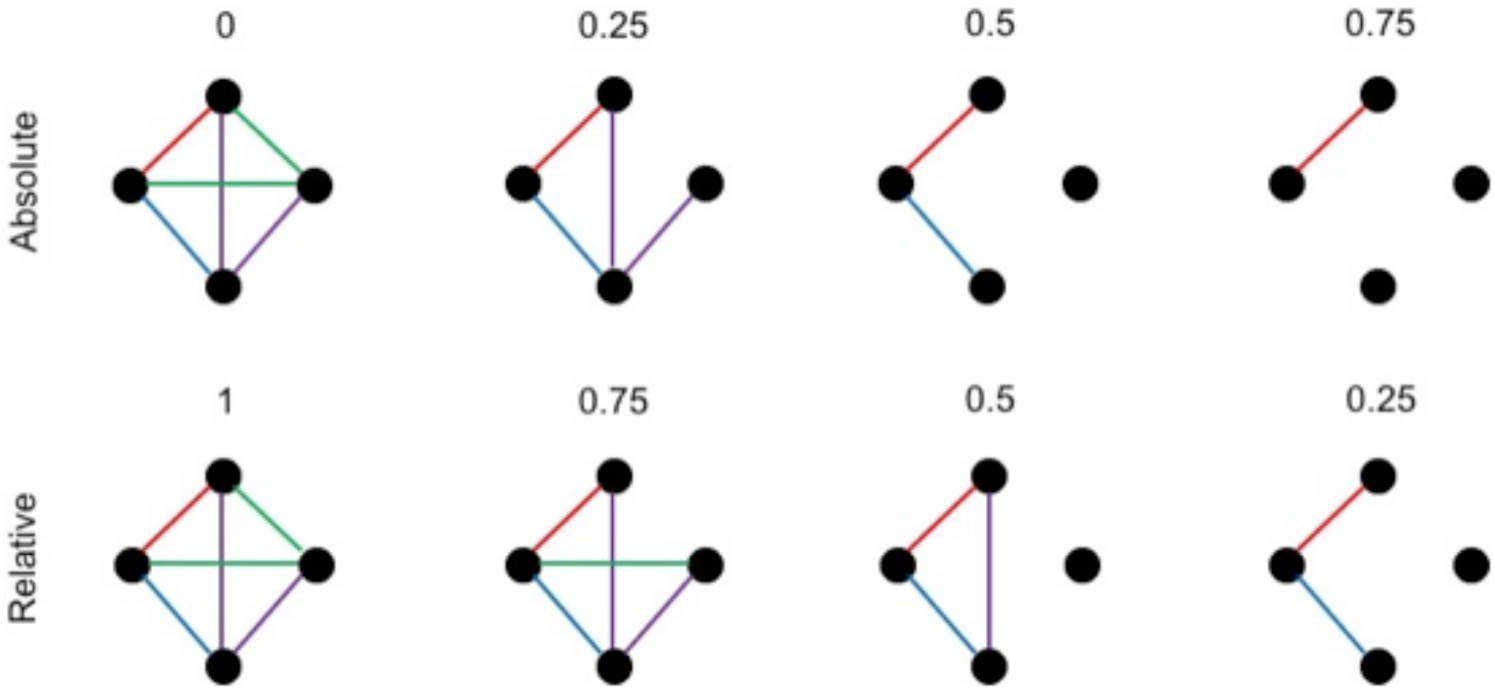
Absolute and Relative Thresholding. Top row: absolute thresholding. With absolute thresholding the connection strengths in the graph are normalized. Subthreshold connections are removed while suprathreshold connections remain. Increasing the thresholds hence sparsifies the graph. Bottom row: relative thresholding. With relative thresholding the threshold indicates the proportion of strongest connections that remains in the graph. Reducing the thresholds hence sparsifies the graph. The colors of the connections indicate their connection strengths. From strongest to weakest: red – blue – purple –green.

**Figure 2.**
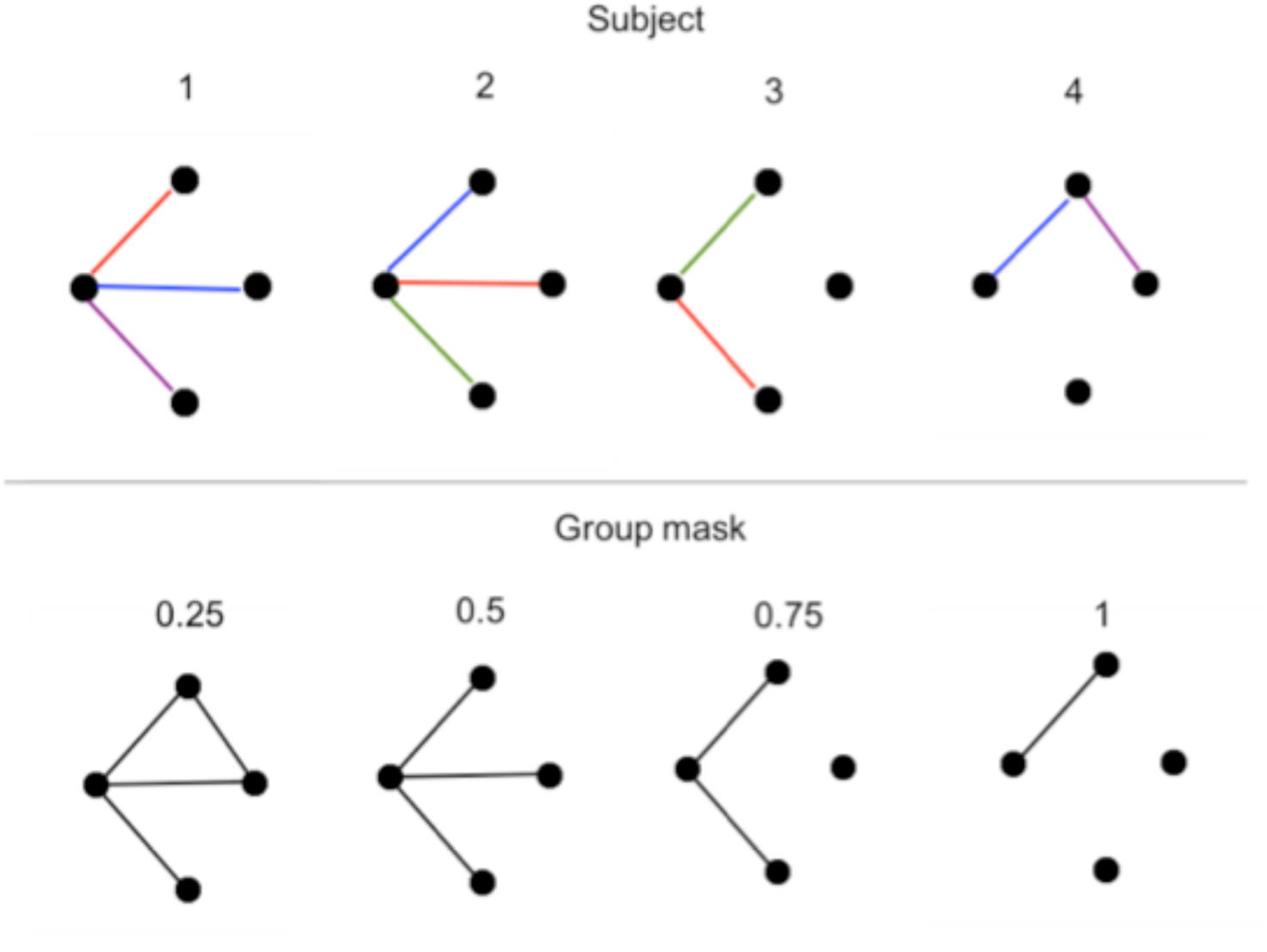
Consensus Thresholding. The top row depicts four fabricated example graphs, where the colors indicate different connection strengths. The group threshold value indicates the proportion of subjects that need to show a connection, regardless of strength, for that connection to be included in the group mask. Increasing the threshold hence sparsifies the mask. Every subject’s connectivity matrix subsequently gets multiplied by the group mask.

While thresholding methods are widely used in the field of structural connectomics one major limitation is the arbitrary nature of threshold constraints and their lack of relevance to the underlying white (matter) biology and inherent biases prevalent in tractography algorithms. A recent major advancement in this regard has been the development of streamline filtering algorithms. The first of these (spherical deconvolution informed filtering of tractograms) SIFT (Smith et al., 2013) was developed in order to address the following issues: i) Longer white matter pathways are present in greater volume and are therefore over defined by streamline reconstruction, ii) streamline tendency to follow the straightest path and, iii) streamlines do not have a volume associated with them in streamline tractography. The SIFT algorithm uses the results of the spherical deconvolution of the diffusion signal to determine which streamlines to remove from the dataset. This results in reconstructed connections, which are proportional to the fiber density as estimated by the diffusion model. As SIFT requires the removal of streamlines this means that the vast amount of streamlines generated are discarded. Therefore connectome reconstruction requires the generation of a large number of streamlines (50-100 million), which requires long processing times and large amounts of memory. While this may be possible when analyzing a few subjects this approach is not feasible for larger datasets. Because of this limitation the successor of SIFT, SIFT2 has been developed more recently. Instead of removing streamlines, in SIFT2 an effective cross-sectional area is determined for each streamline such that the reconstructed fibre volume matches those estimated directly from the diffusion signal (Smith et al., 2015). As this avoids the removal of streamlines, the number required to be generated is in the region of 5-10 million, thus greatly reducing the computation time and memory requirements. However, it is still not clear if these new advancements in streamline filtering will translate into more robust structural connectome construction.

In this study, we thoroughly examine how different thresholding procedures, without and with streamline filtering (based on SIFT2), affect the stability of graph theory metrics, the ability to detect between group differences and correlations with clinical variables. We illustrate these scenarios in a cohort of individuals with clinical and pre-clinical neurodegeneration.

## Materials and Methods

### Participants

A cohort consisting of Huntington’s disease (HD) (*n* = 38), premanifest Huntington’s disease (preHD) (*n* = 50) and control participants (*n* = 47) from the London, Paris and Leiden sites of the TRACK-HD study were included. Structural connectivity changes have been studied previously in this cohort (see (McColgan et al., 2015) for detailed inclusion/exclusion criteria).

### MRI acquisition

T1-and diffusion-weighted images were acquired using Siemens (London and Paris) and Philips (Leiden) 3T MRI scanners. Diffusion-weighted images with 42 unique gradient directions (*b*=1000 sec/mm^2^) were collected with either seven images (Siemens) with no diffusion weighting or one image with no diffusion weighting (Phillips). See (McColgan et al., 2015) for detailed scanning parameters.

### Preprocessing

Cortical and subcortical regions of interest were generated by segmenting a T1-weighted image using Freesurfer (Desikan et al., 2006). These included 70 cortical and six subcortical regions (caudate, putamen and thalamus bilaterally). These targets were warped into diffusion space by finding the mapping between the T1-weighted image and fractional anisotropy map using the NiftyReg toolkit (Modat et al., 2010) and applying the resulting warp to each of the regions of interest. The Freesurfer segmentation was also used to generate foreground masks for the tractography. The graph theoretic analysis uses a foreground mask generated by combining the cortical/subcortical grey matter masks with the white matter mask.

The diffusion data was preprocessed by first generating a brain mask using FSL’s brain extraction tool with b = 0 image (Smith, 2002). This mask was then eroded by one voxel to provide a more stringent mask. Next, eddy current correction was used to align the diffusion-weighted volumes to the first b = 0 image and the gradient directions were updated to reflect the changes to the image orientations. Finally, data were reconstructed using diffusion tensor imaging and constrained spherical deconvolution (CSD), as implemented in MRtrix (Tournier et al., 2012). The CSD reconstruction used a maximum spherical harmonic order of 6 for both the response and the fiber orientation distribution functions.

### Diffusion Tractography

Whole brain probabilistic tractography was performed using the iFOD2 algorithm in MRtrix (Tournier et al., 2012). Specifically, 5 million streamlines were seeded throughout the white matter, in all foreground voxels where fractional anisotropy > 0.2. Streamlines were terminated when they either reached the cortical or subcortical grey matter mask or exited the foreground mask. The reconstructed data was then further processed using SIFT2 to reduce the biases in the reconstructed data (Smith et al., 2015). The resulting set of streamlines and weights were used to construct the structural brain network.

### Streamline filtering

The SIFT algorithm selectively removes individual streamlines from the reconstruction such that the density of the reconstructed connections is proportional to the fiber density as estimated by the diffusion model (Smith et al., 2013). To test the accuracy of the streamline densities, SIFT offers a mechanism to test the quality of the streamlines against the fiber orientation distribution (FOD) lobe integrals, which result from the diffusion data. The algorithm tries to minimize cost function *f*:

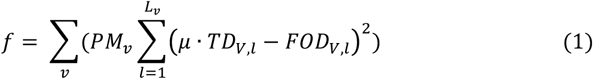

where *v* refers to the voxel and *l* to the lobe number. *PM*_*v*_ is the processing mask in the voxel where the lobe is located, *TD*_*V,l*_ the track density, which is the sum of all the streamlines in the FOD lobe, and *μ* the scaling factor between the units of TD and FOD. When streamlines are removed, the value of *TD*_*V,l*_ reduces and *f* decreases (assuming *TD*_*V,l*_ > *FOD*_*V,l*_). The removal of streamlines is done through ‘gradient descent’ optimization, which seeks to decrease the value of the cost function (Smith et al., 2013).

An alternative method to the removal of streamlines by SIFT is SIFT2, where each streamline gets weighted by determining an appropriate cross-sectional area multiplier. SIFT2 therefore utilizes all streamlines in the reconstruction (Smith et al., 2015). Where SIFT defines the track density (TD) merely as the sum of all streamlines through the FOD lobe, SIFT2 weights each individual streamline by weighting factor 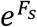, where *F*_*s*_ refers to the weighting coefficient *F* in streamline *s*, before summing them to determine TD. SIFT2 tries to find a vector of weighting coefficients *F* such that when the contribution of each streamline is weighted according to its value in this vector, the streamline densities match the FOD lobe integrals. The cost function that needs to be minimized in SIFT2 is:

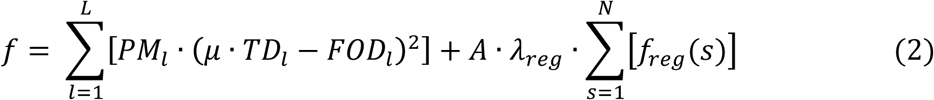

The difference between the cost function used here and the cost function in (1), is the weighting of *TD*_*l*_ and the additional regularization term *f*_*reg*_ (*λ*_*reg*_ is a user-controllable parameter and *A* a scaling constant). *f*_*reg*_ linearly depends on Γ^2^, which is a function of the streamline weighting coefficient *F*_*s*_ and the mean weighting coefficient in the lobe 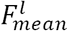:

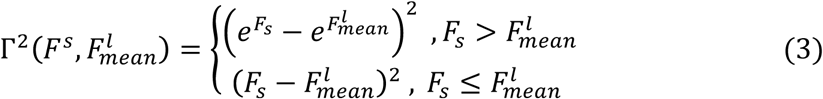

(3) Shows that if the weighting coefficient of the streamline is higher than the mean weighting coefficient, the penalization is directly applied to the weighting factor 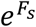. If the weighting coefficient is lower than the mean, the penalty is applied to the weighting coefficients themselves. In this way, streamlines with very large weights are more heavily penalized than streamlines with low weights. This prevents individual streamlines from taking very large weighting factors and obscuring the tractogram.

All analysis in this study were performed without and with streamline filtering using SIFT2 in order to assess the effect of streamline filtering on connectome reconstruction in the context of graph thresholding. The original SIFT algorithm was not used as construction of connectomes with comparable streamlines to SIFT2 would require unfeasible processing times and memory in a cohort of the size used in this study.

### Structural network construction

Regions of interest were defined as connected if a fibre originated in region of interest (ROI) 1 and terminated in ROI 2. The connections were weighted and together formed a 76 × 76, undirected and weighted, structural connectivity adjacency matrix (McColgan et al., 2015). These connectivity matrices were then subjected to relative, consensus and consistency thresholds.

### Relative thresholding

Relative thresholding is based on connection weight, where a proportion of the strongest connections are retained in the connectivity matrix. This allows for consistent network densities across subjects and thus more meaningful group comparisons, however absolute differences in connectivity may be lost (Garrison et al., 2015). In practice relative thresholding is commonly performed over a range of thresholds (Zhang et al., 2011). Once this has been performed a particular threshold level maybe chosen. This may be based on predefined criteria such as the existence of small worldness within a threshold range (Gargouri et al., 2016). An alternative approach is to compute graph theory measures across a range of thresholds and calculate the area under the curve across this range (Drakesmith et al., 2015; Hosseini et al., 2012). In this study the connectivity matrices were thresholded, from dense to sparse, between *t* = 1-0.01 at 0.01 intervals. See Figure 1 for an illustration.

### Absolute thresholding

When using absolute thresholding connections are retained above a specified value and removed below a specified value. This can result in connectomes with different densities across participants, which can pose a problem for between group analyses. This method is most commonly used in functional connectivity, whereby only correlations above a certain value are retained. In this study the connectivity matrices were individually normalized by the largest connection in the matrix and then thresholded, from dense to sparse, with cut-off values ranging between *t* = 0.01-1 at 0.01 intervals. Suprathreshold connections remained in the connectivity matrix and subthreshold connections were removed. Please see Figure 1 (top row) for illustration.

### Consensus thresholding

Consensus thresholding is commonly used in structural connectivity analyses (van den Heuvel and Sporns, 2011). This may be performed across all participants in a study (van den Heuvel et al., 2013) or at the group level (McColgan et al., 2015). While the connections common to a percentage of the group are retained there is the possibility that newly formed connections relating to pathology or compensatory mechanisms in disease may be removed. In this study we created a group thresholding mask based on the control group, as absent connections in the HD groups may be lost due to pathology. A connection remained in the connectivity matrices if it was present in a proportion of the control group. The proportions ranged, from dense to sparse, between *t* = 1-0.01 at 0.01 intervals. See Figure 2 for an illustration of this thresholding scheme.

### Consistency thresholding

Structural connectome studies are increasingly adopting probabilistic tractography as this approach is more effective at resolving crossing fibres compared to deterministic approaches. One consequence of this is the creation of denser connectomes. This poses problems for consensus thresholding as networks are nearly fully connected in all participants. While relative thresholding is one alternative, thresholding based on connection weight can introduce bias to the connectome. It is shown that weight based relative thresholding penalises long range connections, as they tend to have weaker connection strength when compared to short range connections (Roberts et al., 2016). In order to address this issue (Roberts et al., 2016) proposed a novel consistency based thresholding method such that connections are ranked based on their coefficient of variation across a group of participants. Connections with the lowest coefficient of variation are then retained and those with a high coefficient of variation are discarded. In this study we perform consistency thresholding on the healthy control group and used the resulting binarised matrix to remove connections at the subject level for all participants. The proportions ranged, from dense to sparse, between *t* = 1-0.01 at 0.01 intervals.

### Graph theoretical analysis

In computational neuroscience, graphs can be used to represent anatomical (or functional) connections between brain areas that interact to give rise to various cognitive processes, where the vertices represent different areas of the brain and the edges represent the connections between those areas (Bullmore and Sporns, 2009). Graph theoretic descriptions have been very useful in terms of defining neurological disorders as disconnectivity of the putative graph made up of the interacting populations of neurons (Schroeter et al., 2015). By using graph theoretic descriptions one can summarise different aspects of the large neural network’s structure and topology. For doing this different network metrics or descriptors are in use, which we now summarise.

### Network metrics

Network metrics are statistical measures that characterize certain aspects of a network. These can be divided into local metrics that define brain network topological characteristics at the regional level or global metrics that define the overall functioning of the network as a whole. Metrics may be binary in that they are derived based on the presence or absence of connections or weighted whereby they take into account the magnitude of connection strength. This may either be based on streamline counts or fractional anisotropy (FA) in the case of structural connections or temporal correlations of fMRI time series between regions in the case of functional connectomes or directed effective connectivity (Hyett et. al JAMA, Psychiatry 2015) as measured by dynamic causal modelling (Razi & Friston, 2016). However, in this work we focus only on structural connectomes. From each connectivity matrix, four network metrics were calculated: degree, strength, modularity and global efficiency. The network metrics were calculated using the brain connectivity toolbox (Rubinov & Sporns, 2010) and their graphical illustrations are shown in Figure 3.

**Figure 3.**
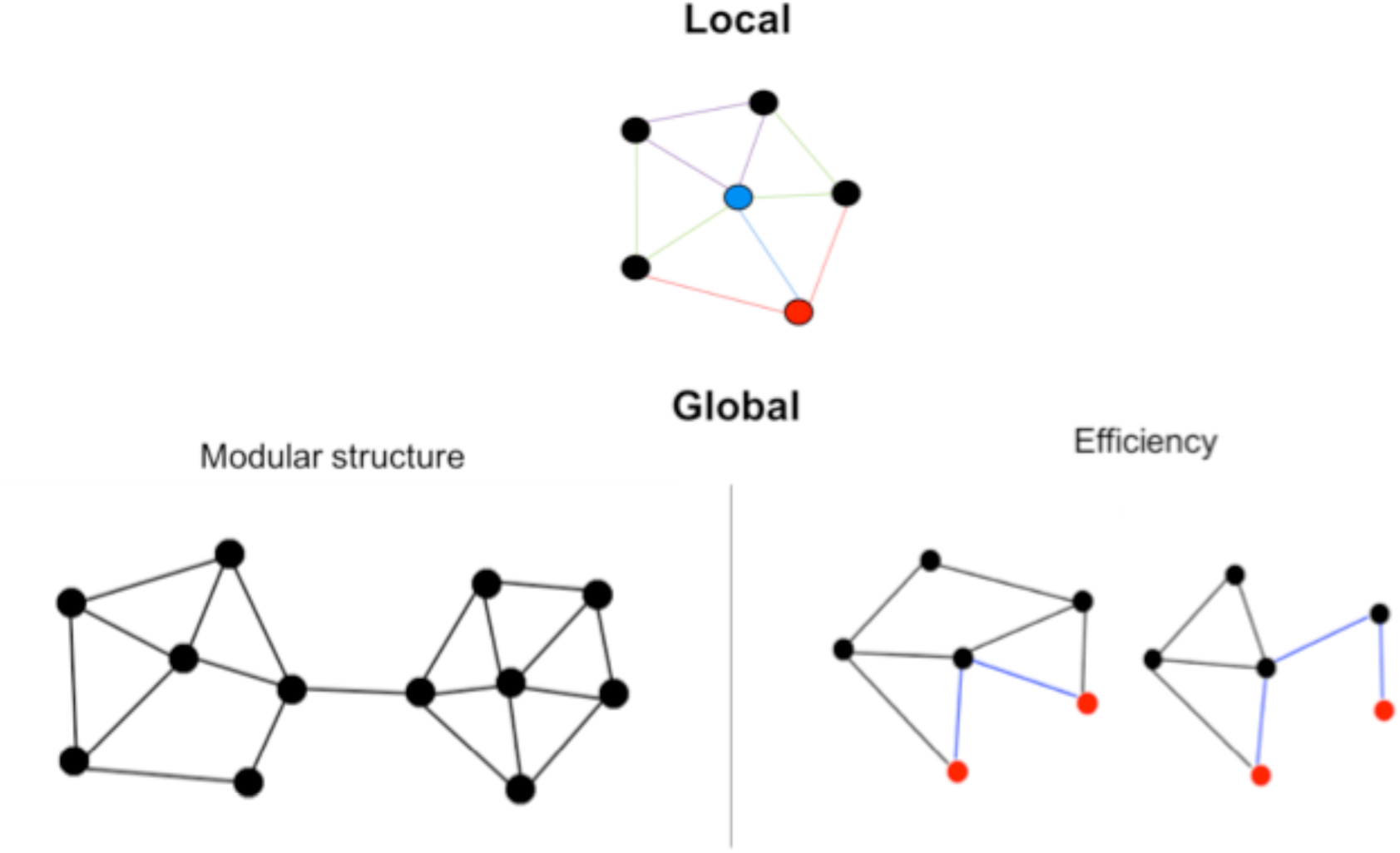
Local Graph Metrics: the degree of a node is the number of edges connected to that node. In this graph the blue node therefore has a higher degree (k = 5) than the red node (k = 3). The strength of a node is the sum of the connection weights of all edges connected to that node. The colors of the connections represent the connection strengths with red = 4, blue = 3, purple = 2 and green = 1. The red node has a higher strength (S = 11) than the blue node (S = 9). Global: the modularity of the network is a measure of segregation. A modular structure has segregated clusters of nodes with a high number of within-connections and a low number of between-connections. The global efficiency of the network is a measure of integration. Global efficiency is defined as the inverse of the shortest path length between two nodes. The efficiency between the two red nodes in the left graph is higher than in the right graph.

### Local metrics

Local metrics often quantify connectivity profiles of individual nodes and therefore reflect the way node is embedded in the network (Rubinov and Sporns, 2010). This allows us to make inferences about functions of specific brain regions and their role in disease processes. The most fundamental metric provided by a graph is *degree*. The degree of a node is equal to the number of edges connecting that node to the rest of the network. An additional regional metrics is *strength*. Strength is defined as the sum of all connection weights connected to that node (see Figure 3). Degree and strength give insight into the importance of nodes in the network.

### Global metrics

Global metrics can inform us about the brain network at a systems level. They can characterize both the integration and the segregation of the network. Metrics of structural segregation quantify the presence of clusters or modules in the network. The presence of such clusters or modules suggests the potential for functional segregation. High segregation indicates the network’s ability for specialized processing. An important measure of segregation is *modularity*, which represents the modular structure within the brain network. Modularity can be defined as the degree to which a network can be subdivided into groups of nodes that have a large number of within-connections and fewer number of between-connections. See an illustration of a modular structure of a graph in Figure 3.

Metrics of structural integration represent the brain’s ability to rapidly combine specialized information from distributed brain regions. For example the *path length* between two nodes estimates the potential for those two nodes to integrate information: the shorter the path length the higher the potential for integration (see Figure 3). *Global efficiency* is the average of the local efficiencies for individual nodes. Mathematically efficiency and path length are inversely proportional. However, efficiency is a preferred metric compared to the path length since in a disconnected network, the path length between two nodes residing in two unconnected modules will be infinite whereas the efficiency will be zero which is numerically easier to deal with in the graph theoretic analysis (Rubinov and Sporns, 2010).

### Network Stability

Network stability was defined by how stable the graph theory metric values were over a range of thresholds. Each network measure was calculated for every threshold value and a ‘survival curve’ was then plotted in order to visualize the stability of the network measures across thresholds (Garrison et al., 2015). This survival curve describes how the network measure changes depending on the threshold applied. Since the network metric can be estimated by a single point on the curve, the shape of the curve indicates the stability of the network metric across thresholds (Garrison et al., 2015). For investigating the stability of group differences for different thresholding schemes across various graph network metrics we used the changes in the direction of group differences to indicate instability. Depending on the threshold used, the group difference can be reversed hence providing a measure of network stability. To test for group differences permutation testing, using unpaired *t*-tests, was performed with 10,000 permutations at each threshold (Nichols and Holmes, 2002). A group difference was deemed significant if *p* < 0.05. We then computed the direction of statistical significance in group differences for each metric and thresholding method using left and right one-tailed t-tests.

### Clinical correlations

To test which thresholding method is the most appropriate to investigate structural connectivity loss in neurodegenerative disease, we correlated the strength of the right caudate with clinical measures over the whole range of thresholds for preHD and HD groups combined. We selected right caudate because the caudate is one of the first structures to be affected in HD (Tabrizi et al., 2011). The clinical measures used were Disease Burden Score (DBS) and Negative Emotion Recognition. DBS is computed based on the participants’ age and CAG repeat (Penney et al., 1997). Emotion recognition was chosen due to its sensitivity in Huntington’s disease as previously reported (McColgan et al., 2015; Novak et al., 2012). We used Pearson correlation to control for age, sex, study site, education and CAG repeat.

## Results

### Graph Metrics

### Degree

We compared the degree and the strength of the right caudate for various thresholding approaches due to the significance of this brain region in Huntington’s disease as caudate showed both volumetric and structural connectivity loss in the premanifest stage (McColgan et al., 2015; Tabrizi et al., 2011). Figure 4 summarises the results for local metric of degree in which we plotted survival curves for all of the four thresholding strategies (rows) with and without SIFT2 (columns). Each panel of Figure 4 has three survival curves for manifest HD (blue), premanifest HD (green) and healthy controls (red). The thresholds on the x-axis are such that the network becomes dense to sparse going from left to right. For both relative and consistency thresholding there was an almost linear decline in degree as threshold level increased (from dense to sparse). This is unsurprising as the former method retains a proportion of connections based on weights while the latter retains connections based on coefficient of variation. For absolute thresholding, the degree drops sharply at low threshold levels (that corresponds to denser networks). Consensus thresholding was stable across the large range of thresholds and starts to decrease only at very high threshold levels corresponding to very sparse networks. This is likely due to the use of dense connectivity matrices created when using probabilistic tractography. Qualitatively between group differences are most visible for consensus thresholding. Very little difference is seen without and with the application of streamline filtering using SIFT2. This may be due to the fact that SIFT2 weights connections as opposed to removing them and so would have very little effect on a binary metric such as degree.

**Figure 4.**
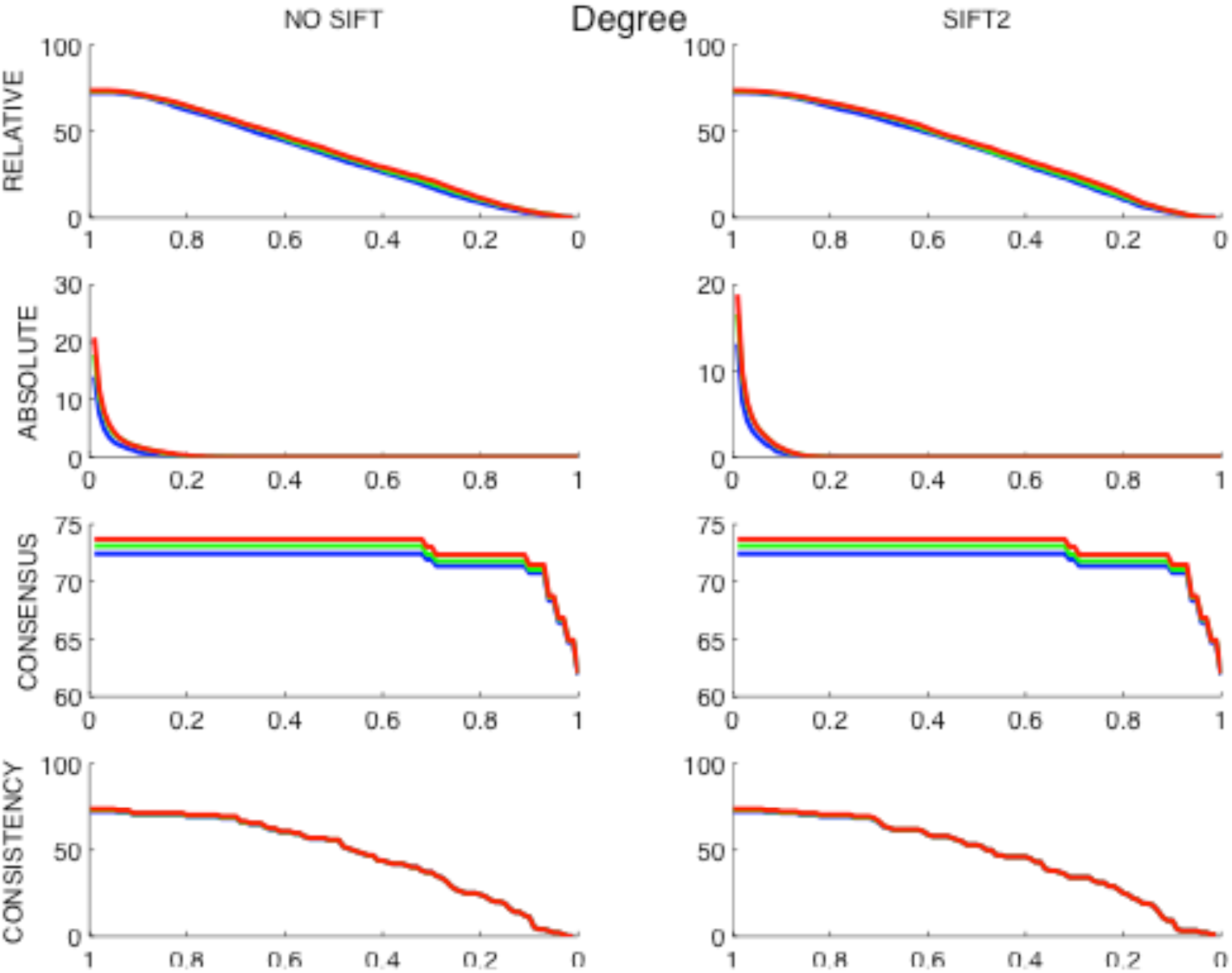
Degree of right caudate across thresholds. The left columns show the metrics for no streamline filtering and the right columns for streamline filtering (SIFT2). The blue survival curve shows the metrics for manifest Huntington’s disease participants, the green survival curve for premanifest Huntington’s disease participants and the red survival curve for control participants. The X-axes depict the threshold level, with proportion of connections included in the network for relative thresholding, cut-off value for absolute thresholding and proportion of connections included in the mask for consensus and consistency thresholding. The Y-axes depict the degree.

### Strength

Figure 5 shows the survival curves for the graph metric of strength using different thresholding strategies which is in the same format as the previous figure. When compared with degree, the strength metric shows greater stability for both relative and consistency thresholding. Stability is maintained up to 0.4 and values begin to drop thereafter. For strength, absolute thresholding displays a similar profile as it does with the degree metric, with a sharp drop in metric value at low thresholds (more dense network). The strength metric is largely unchanged by consensus thresholding, which is a likely consequence of performing this approach on almost fully connected matrices. Qualitatively group differences are visible for all threshold approaches, suggesting in the context of densely connected matrices generated using probabilistic tractography, strength is a more appropriate regional network metric than degree. The application of SIFT2 streamline filtering results in lower strength metric values for all thresholding approaches. For consistency thresholding streamline filtering seems to increase the stability of strength across thresholds when compared to no streamline filtering.

**Figure 5.**
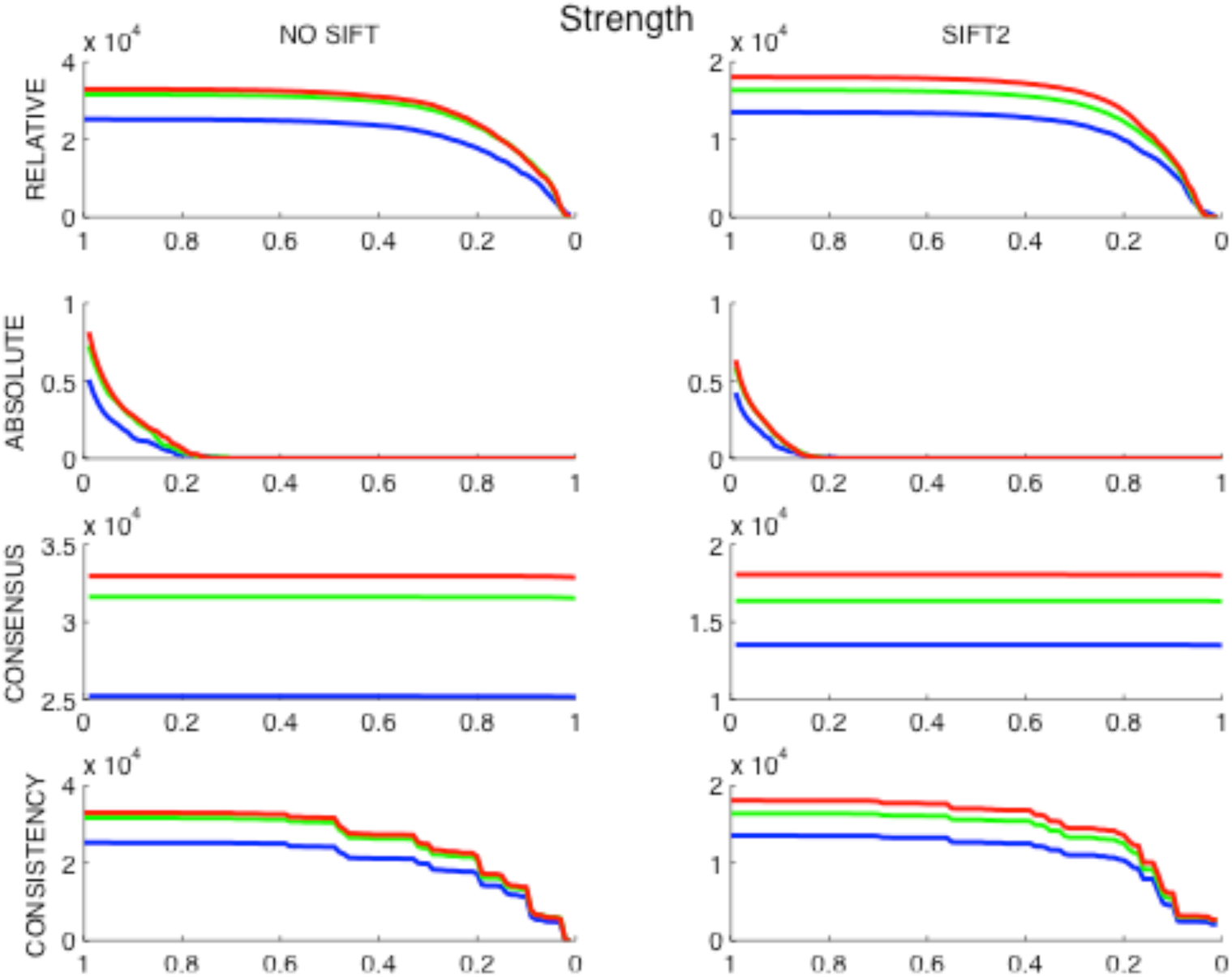
Strength of right caudate across thresholds. The left columns show the metrics for no streamline filtering and the right columns for streamline filtering (SIFT2). The blue survival curve shows the metrics for manifest Huntington’s disease participants, the green survival curve for premanifest Huntington’s disease participants and the red survival curve for control participants. The X-axes depict the threshold level, with proportion of connections included in the network for relative thresholding, cut-off value for absolute thresholding and proportion of connections included in the mask for consensus and consistency thresholding. The Y-axes depict the degree or strength value respectively.

### Modularity

Figure 6 summarises the results for modularity as the metric of segregation for various thresholding strategies in the same format as in the previous two figures. For relative and consistency thresholding, modularity increases as the network becomes spraser due to the removal of connections hence increasing the segregation of the network. Absolute thresholding is more stable for modularity than for the local metrics of degree and strength, with modularity dropping at high threshold values i.e. when the network becomes sparser. Consensus thresholding has minimal effect on the modularity metric. Qualitatively group differences are most visible for consensus thresholding. With the application of streamline filtering group differences are more visible for consensus thresholding.

**Figure 6.**
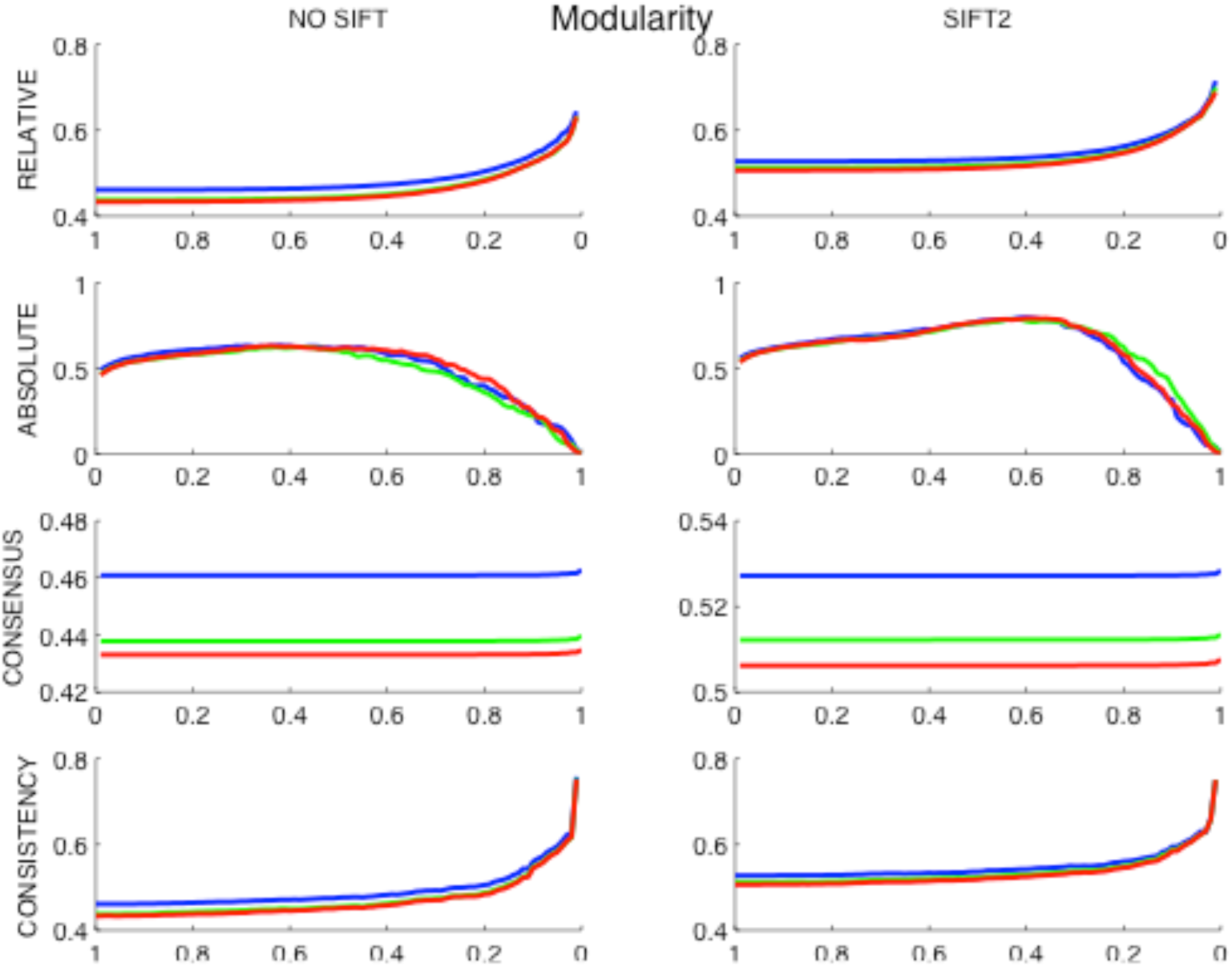
Modularity across thresholds. The left columns show the metrics for no streamline filtering and the right columns for streamline filtering (SIFT2). The blue survival curve shows the metrics for manifest Huntington’s disease participants, the green survival curve for premanifest Huntington’s disease participants and the red survival curve for control participants. The X-axes depict the threshold level, with proportion of connections included in the network for relative thresholding, cut-off value for absolute thresholding and proportion of connections included in the mask for consensus and consistency thresholding. The Y-axes depict modularity.

### Global Efficiency

Figure 7 summarises the results for global efficiency as metric of network integration. Global efficiency is perhaps the most stable metric for relative and consistency thresholding with values only dropping at very high thresholds when the network becomes very sparse. In comparison, absolute thresholding shows an almost linear decrease. As with other network metrics in previous figures, global efficiency is largely unaffected by consensus based thresholding and is the most stable. Streamline filtering has a dramatic impact on qualitative group differences for consensus thresholding, without filtering the manifest HD group has a higher global efficiency than controls, however this is reversed when streamline filtering is applied which is a very interesting observation. SIFT2 has no clearly visible impact on the remaining thresholding strategies.

**Figure 7.**
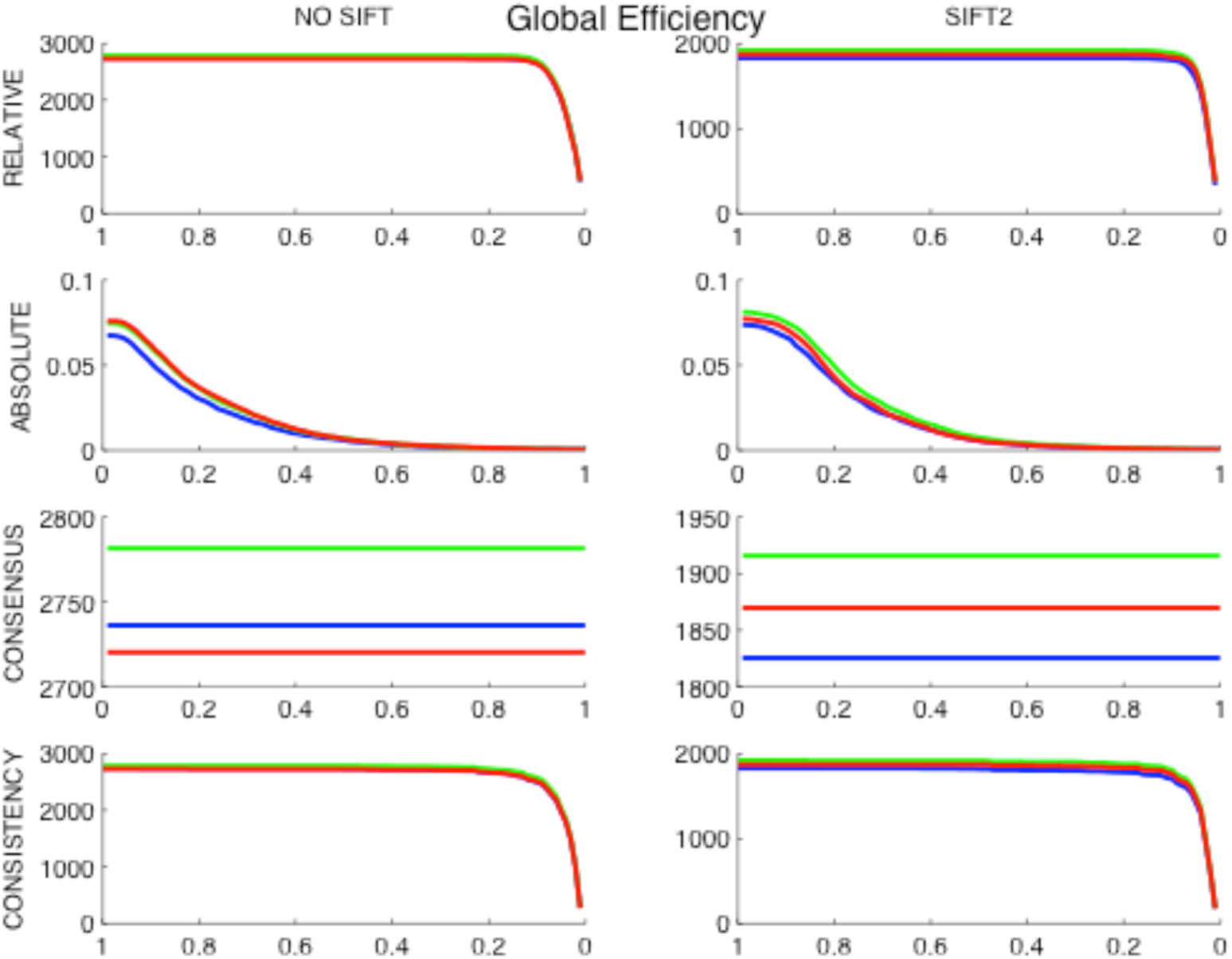
Global efficiency across thresholds. The left columns show the metrics for no streamline filtering and the right columns for streamline filtering (SIFT2). The blue survival curve shows the metrics for manifest Huntington’s disease participants, the green survival curve for premanifest Huntington’s disease participants and the red survival curve for control participants. The X-axes depict the threshold level, with proportion of connections included in the network for relative thresholding, cut-off value for absolute thresholding and proportion of connections included in the mask for consensus and consistency thresholding. The Y-axes depict global efficiency.

### Group Differences

Group differences indicate a significant difference in network metric between i) manifest HD and controls, ii) manifest HD and preHD and, iii) preHD and controls groups. To statistically test for group differences, permutation testing, using unpaired *t*-tests, was performed with 10,000 permutations at each threshold (Nichols and Holmes, 2002). A group difference was deemed significant if *p* < 0.05. The outcome of these statistical tests for group differences are summarized in Table 1.

**Table 1.**
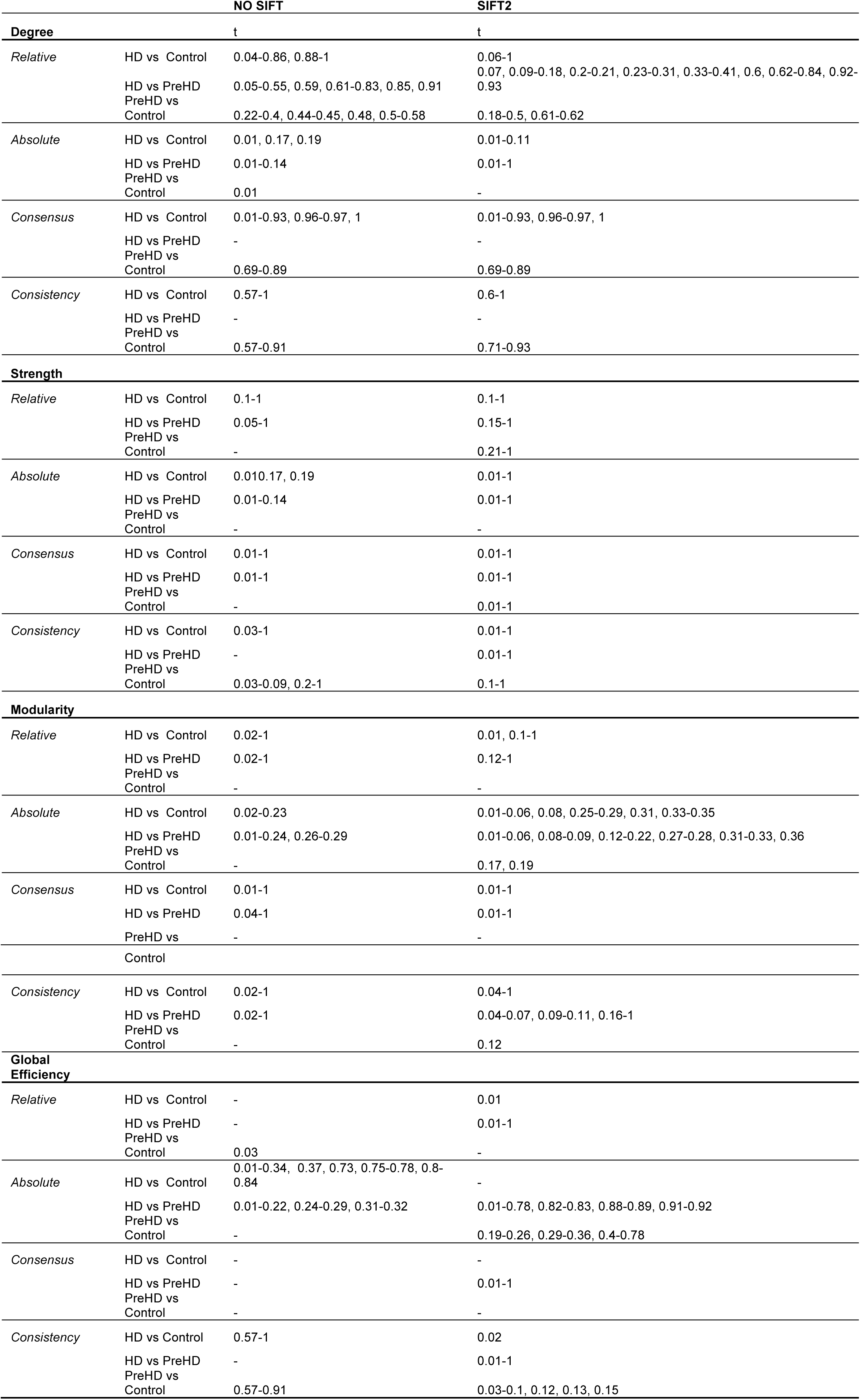
Significant Group Differences across thresholds. Group differences are deemed significant if P < 0.05 two-tailed. t = threshold, HD = manifest Huntington’s Disease group, preHD = premanifest Huntington’s Disease group.

For degree, group differences are seen between all groups for relative thresholding however this varies across thresholds. For both consensus and consistency thresholding no group differences are seen between HD and preHD. For absolute thresholding, group differences are seen across all groups when the network is very dense.

For the strength metric, group differences are more stable across thresholds. The application of SIFT2 streamline filtering results in the detection of group differences between preHD and controls for both consensus and consistency thresholding that were not present without streamline filtering. PreHD vs. control group differences were absent for absolute thresholding both with and without SIFT2.

For modularity, group differences varied across thresholds. PreHD vs. control group differences were generally absent apart from absolute and consistency thresholding after the application of streamline filtering. For global efficiency group differences were generally absent for relative and consensus thresholding prior to streamline filtering. For absolute and consistency thresholding group differences varied across thresholds.

### Direction of group differences

To investigate whether the direction of significant group differences were consistent across thresholds, left and right one-tailed t-test were done and p-values were calculated for each graph theory metric for every thresholding strategy across thresholds. The results are summarised in Table 1 where total number of significant group difference, at different thresholds, is calculated for various thresholding strategies.

Directions of group differences were consistent for both degree and strength for all four thresholding approaches. The application of streamline filtering altered the number of thresholds where group differences were seen but not the direction of significance. For strength, streamline filtering resulted in the consistent detection of group differences between preHD vs. controls across a large number of thresholds for relative, consensus and consistency thresholding.

For modularity relative, consensus and consistency thresholding showed consistently right sided group differences. For absolute thresholding, the group differences were seen for both left and right sided t-tests depending on the chosen threshold hence was less consistent.

For global efficiency group differences between HD vs. controls were seen for only a few thresholds. For relative thresholding right-sided group differences were seen without streamline filtering at one threshold, while left-sided group differences were seen at 4 thresholds with SIFT2 streamline filtering. For absolute thresholding left sided group differences were seen without and with SIFT2 streamline filtering. Consensus and consistency thresholding showed consistent directions of group differences.

### Clinical Correlations

To investigate the effect of different thresholding approaches and the application of streamline filtering on clinical correlations, we preformed partial correlations analysis between strength of the right caudate for HD gene carriers and disease burden score (DBS) (see Figure 8) and strength of the right caudate and emotion recognition (see Figure 9) while controlling for age, gender, site, education and CAG repeat. For DBS, in Figure 8, a negative correlation was seen with strength of the right caudate. This finding was stable across thresholds for relative and consensus thresholding. There was some variability across consistency thresholds. Absolute thresholding was very unstable across threshold values. The application of streamline filtering increased the strength of correlation for consistency thresholding.

**Figure 8.**
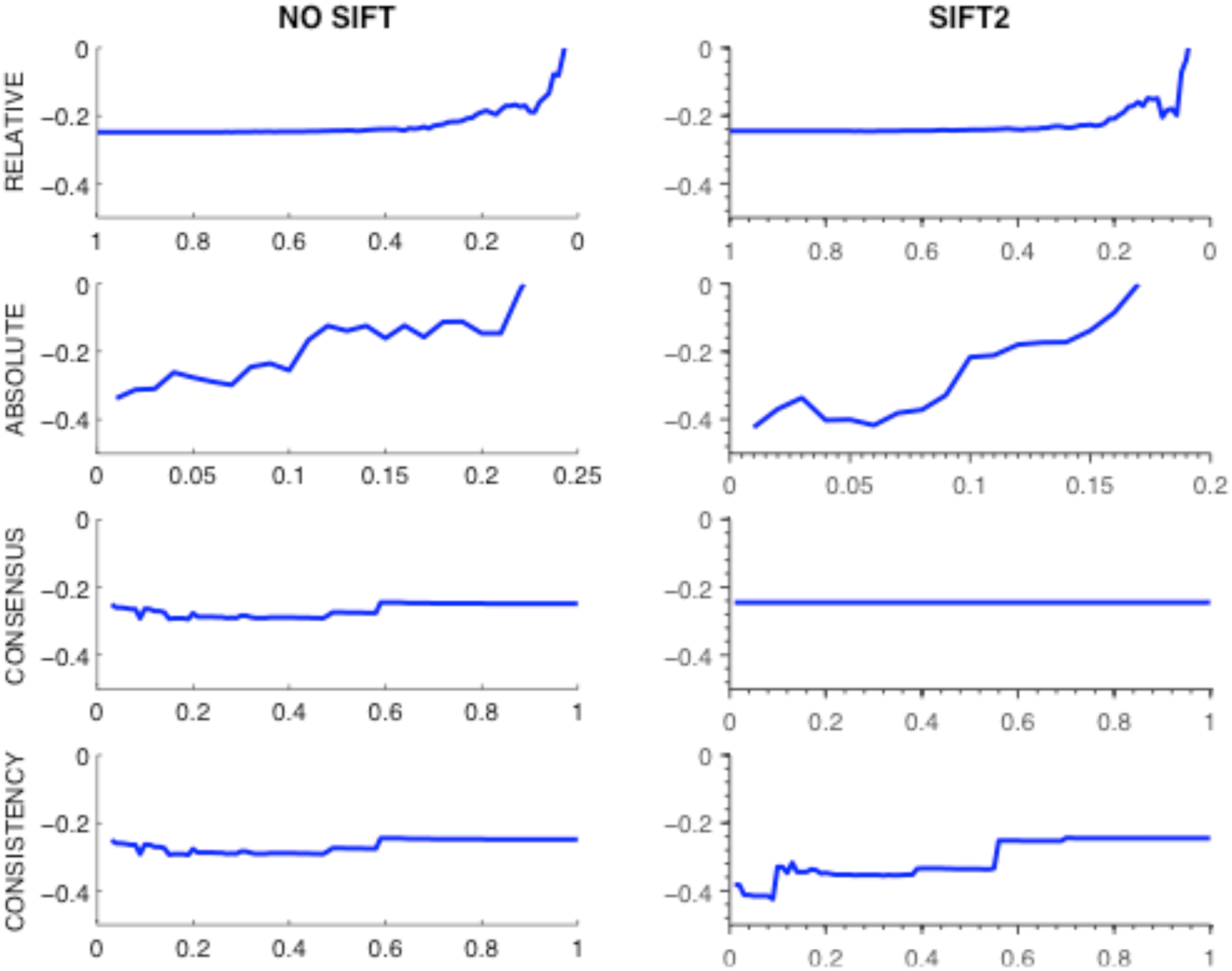
Correlation between strength of the right caudate and disease burden. The left columns show the correlations for streamline filtering and the right columns for SIFT2. The X-axes depict the threshold level, with proportion of connections included in the network for relative thresholding, cut-off value for absolute thresholding and proportion of connections included in the mask for consensus and consistency thresholding. The Y-axes depict the correlation coefficient. The correlation with disease burden is computed for manifest and premanifest Huntington’s disease groups.

**Figure 9.**
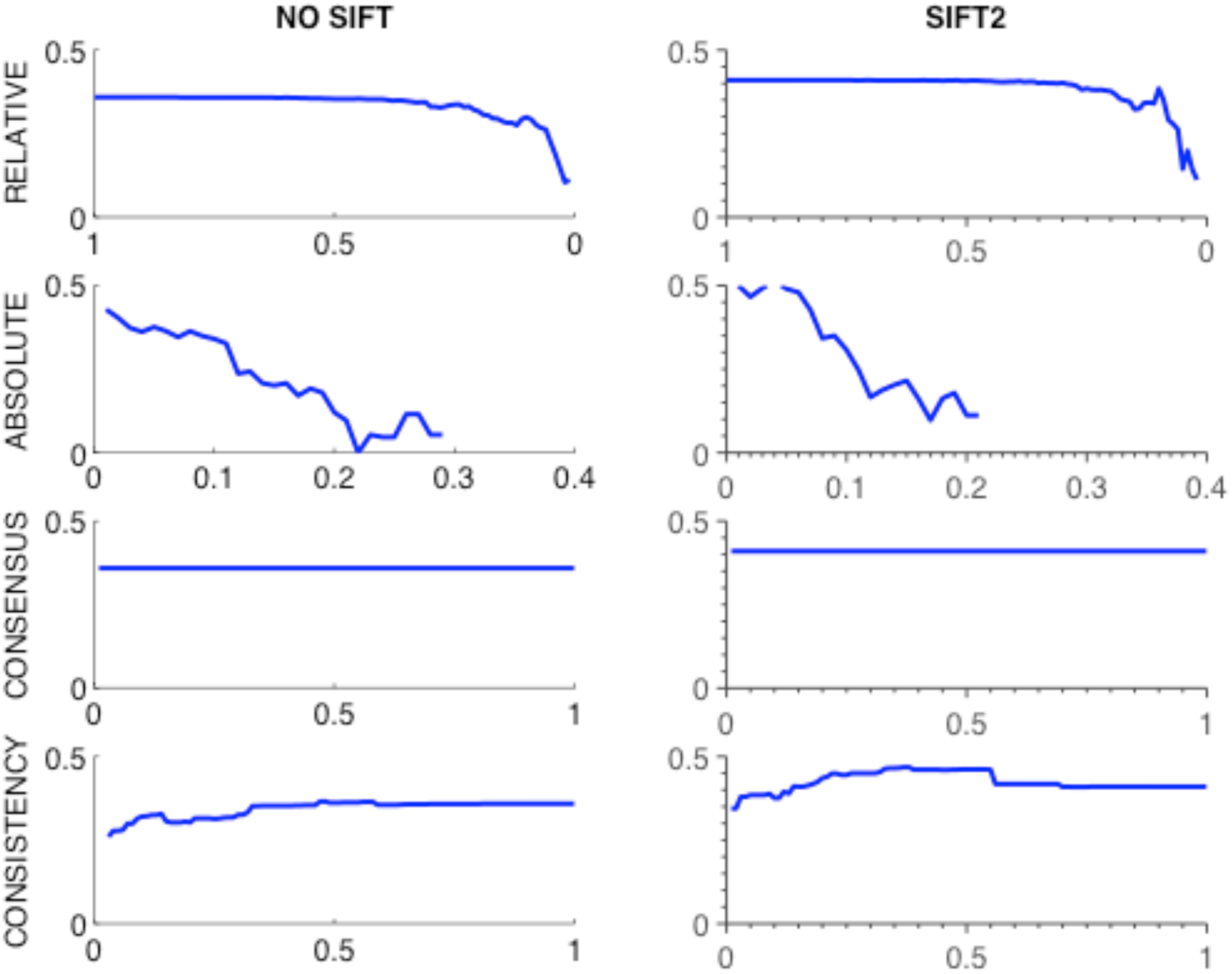
Correlation between strength of the right caudate and emotion recognition. The left columns show the correlations for streamline filtering and the right columns for SIFT2. The X-axes depict the threshold level, with proportion of connections included in the network for relative thresholding, cut-off value for absolute thresholding and proportion of connections included in the mask for consensus and consistency thresholding. The Y-axes depict the correlation coefficient. The correlation with emotion recognition is computed for manifest and premanifest Huntington’s disease.

Positive correlations were seen between strength of the right caudate and emotion recognition score. This was stable across thresholds for relative, consensus and consistency thresholding but highly variable for absolute thresholding. For consensus and consistency thresholding the application of SIFT2 streamline filtering increased the strength of correlation.

## Discussion

The aim of this study was to examine the effect of various graph thresholding approaches and streamline filtering algorithm on the stability of structural graph theory metrics evidenced in cohorts with preclinical and clinical neurodegeneration. The four thresholding approaches compared were relative, absolute, consensus and consistency thresholding. For relative, consensus and consistency thresholding, metrics were generally stable across thresholds, apart from very high threshold values when the connectivity matrix becomes very sparse. For all thresholding approaches the presence of significant group differences was variable based on the threshold value chosen. This was particularly the case for degree and relative thresholding and for global metrics (modularity and global efficiency) and absolute and consistency thresholding. The direction of significant group differences was generally consistent apart from absolute thresholding and modularity.

We also assessed the effect of SIFT2 streamline filtering on graph theoretic metrics. SIFT2 had very little effect on the graph theory metric survival curves, however it did result in changes in the absolute value of metrics when compared to no filtering. With respect to group differences, with SIFT2 significant differences were detected between preHD vs. controls for strength in the context of relative, consensus and consistency thresholding. This suggests that SIFT2 may increase the sensitivity to detecting group differences particularly in the premanifest cohort compared to controls. SIFT2 also resulted in higher correlations with two different clinical variables, in the context of consistency thresholding, for the metric of strength of the right caudate and disease burden score. This raises the possibility that the application of SIFT2 may result in a connectome, which is more representative of the underlying biology of the white matter in a brain network.

Relative thresholding is perhaps one of the more widely used thresholding approaches which is either performed at a predefined threshold level (Zhang et al., 2011) or across a range of thresholds (Hosseini et al., 2012). While we show that relative thresolding is stable across thresholds the variability with respect to significant group differences is concerning and suggests that results may differ depending on the choice of the selected threshold. Absolute thresholding, which is used mainly in functional connectomic studies (Harrington et al., 2015), is found to be unstable and thus an inappropriate approach with respect to structural connectomics. While consensus thresholding is stable across thresholds and results in consistent group differences this thresholding technique has very little effect on connectivity matrices that are fully connected. Consistent thresholding has only recently been developed (Roberts et al., 2016) and this is the first study where it has been applied to a clinical population. While it results in consistent graph theory metrics across thresholds, the detection of significant group differences may vary depending on the threshold chosen. Thus, our findings suggest that no thresholding approach evaluated here is without any caveats.

Prior to the development of consistency thresholding, (Drakesmith et al., 2015) proposed multi-threshold permutation correction. However this technique still requires the testing for group effects across multiple thresholds and as with the approaches investigated here there is no principled way of choosing the appropriate threshold. Our findings suggest that the presence or absence of group differences is dependent on the choice of threshold used. We have also shown that the detection of group differences can vary across thresholds, which poses significant methodological concerns for graph thresholding in general.

One approach to remove false positives prevalent in probabilistic tractography generated connectomes (Zalesky et al., 2016) is streamline filtering. While the uptake of this approach in the neuroimaging community has been slow this is most likely due to the large computation times required in the very first method developed, SIFT (Smith et al., 2014). With the more recent development of SIFT2, dense connectomes can now be created with modest processing times (Smith et al., 2015). In this work we show that following the application of SIFT2, there is an increased ability to detect group differences between preHD and controls. Furthermore, stronger clinical correlations are seen after the application of SIFT2. This suggests that SIFT2 may enable the construction of a connectome that is more representative of the underlying white matter biology. With the results we have presented, it may be that the application of SIFT2 without the use of any graph thresholding may be sufficient for connectome construction.

With respect to the graph theory metrics in the context of dense connectomes, we investigated both degree and strength. Degree represents the number of binary connections to a node or brain region and can be used to investigate the importance of a node in a binary network. However, the connectomes resulting from fibre orientation distribution (FOD)-based tractography, implemented in MRtrix, are highly connected. FOD-based tracking is a subcategory of probabilistic tractography, where the FOD is directly sampled during tracking. A lot of probabilistic tractography algorithms merely sample the uncertainty in the dominant orientation without taking the underlying fibre orientation dispersion into account. However by sampling directly from the FOD, streamlines emanating from a common point tend to disperse from one another more (Tournier et al., 2012). When enough streamlines are sampled, any two brain regions can therefore become connected (Bastiani et al., 2012). This makes a binary measure such as degree an uninformative metric when using this framework. Strength on the other hand sums the weights of all edges connected to a node. Thus it is possibly a more appropriate measure in weighted networks, since it uses all the information available from the connectivity matrix.

One caveat to the conclusions drawn from this study are the preprocessing options used in generating connectomes. The effect of graph thresholding strategies may vary based on image acquisition, number of regions of interest in an atlas parcellation and whether healthy controls or clinical populations are being investigated.

## Conclusions

In this work, we investigated a number of graph thresholding procedures for the construction of structural connectomes using diffusion tractography. We showed that the graph metrics calculated following these thresholding strategies are fairly stable across a range of thresholds. However, more crucially, the detection of group differences was highly variable and depended on the specific amount of threshold chosen. This poses significant methodological concerns for graph thresholding for structural connectome construction in general. We showed that using an approach based on streamline filtering using SIFT2 is more sensitive for detection of group differences and it also provided stronger clinical correlations in the Huntington’s disease cohort we used in this study. Therefore, we argue that the application of SIFT2 without graph thresholding may be sufficient for structural connectome construction for group studies in clinical populations.

## Acknowledgements

This study was funded by the Wellcome Trust (GR, PMC) (091593/Z/10/Z, 515103) and supported by the National Institute for Health Research [NIHR] University College London Hospitals [UCLH] Biomedical Research Centre [BRC]. Track-HD is funded by the CHDI foundation, a not for profit organisation dedicated to finding treatments for Huntington’s disease.

---

**TRACK-HD Investigators:**

**TRACK-HD Investigators:**
|  | **Last name** | **First name** | **Location** |
| --- | --- | --- | --- |
| 1. | Bohlen | Stefan | Muenster |
| 2. | Crawford | Helen | UCL |
| 3. | Hart | Ellen P | LUMC, Leiden |
| 4. | Hobbs | Nicola Z | UCL |
| 5. | Jauffret | Celine | Paris |
| 6. | Francisque | Hélène | Paris |
| 7. | Labuschagne | Izelle | Monash |
| 8. | Lahiri | Nayana | UCL |
| 9. | Lehericy | Stéphane | Paris |
| 10. | Monin | Marie Laure | Paris |
| 11. | Malone | Ian | UCL |
| 12. | O'Regan | Alison | Monash |
| 13. | Queller | Sarah | Indiana |
| 14. | Read | Joy | UCL |
| 15. | Reilmann | Ralf | Meunster/GHI |
| 16. | Rödig | Verena | LUMC, Leiden |
| 17. | Valabrègue | Romain | Paris |
| 18. | van der Grond | Jeroen | LUMC, Leiden |
| 19. | Whitehead | Daisy | UCL |
| 20. | Felix | Ttita | Ulm |
| 21. | Katja | Vitkin | Ulm |
| 22. | Theresia | Kelm | Ulm |

